# Lymphatic vessel dysfunction contributes to severe dengue pathogenesis

**DOI:** 10.64898/2026.03.27.714698

**Authors:** Fatima Abukunna, Daniel Matamala Luengo, Aitana Martin Manrique, Justin Duruanyanwu, Matthew Sherwood, Priyanka Patel, Mark Crabtree, Graeme M. Birdsey, Kevin Maringer, Paola Campagnolo

## Abstract

Dengue virus (DENV) infection is a major global health threat, affecting more than half of the world’s population. Severe dengue is a life-threatening condition characterised by systemic bleeding, vascular leakage, and interstitial fluid accumulation that can progress to hypovolaemic shock. Circulating DENV non-structural protein 1 (NS1) has long been implicated in driving vascular hyperpermeability through its disruptive effects on endothelial cell junctions and the glycocalyx.

The lymphatic system, which runs alongside the vascular network, plays a critical role in resorbing and recirculating interstitial fluid and immune cells extravasated from blood vessels. Despite its importance in maintaining tissue fluid homeostasis, the impact of dengue disease on lymphatic vessels has not previously been explored.

Here, we present the first evidence that DENV-2 NS1 induces marked hyperpermeability in lymphatic endothelial cells, as measured by transendothelial electrical resistance, and impairs lymphangiogenesis in vitro. These effects were not attributable to changes in cell viability, morphology, or metabolic activity, as assessed by live/dead and metabolic assays and image analysis. Instead, we observed a defect in lymphatic endothelial cell migration, measured by scratch assay, which may underlie the reduced lymphangiogenic potential. Bulk RNA-seq, immunocytochemistry, and advanced image analysis further demonstrated pronounced reorganisation of cell–cell junctions, the cytoskeleton, and focal adhesions. Notably, junctional proteins including VE-cadherin, ZO-1, and Claudin-5 were not downregulated but instead displayed disorganised distribution along the cell junctions or aberrant cytoplasmic localisation. These structural disruptions became even more pronounced under flow conditions produced using a microfluidic system.

Together, these findings demonstrate for the first time that DENV-2 NS1 directly disrupts lymphatic endothelial cell function, leading to junctional disorganisation and hyperpermeability. Such impairment of lymphatic drainage may contribute to the pathophysiology of severe dengue.

**Author Summary:** Dengue is a rapidly expanding mosquito-borne disease that now affects many tropical and subtropical regions worldwide. Severe cases can lead to extensive fluid leakage from blood vessels, which causes tissue swelling and, in the most dangerous situations, shock. Although much research has focused on how dengue damages the blood vascular system, almost nothing is known about its impact on the lymphatic system, which is responsible for removing fluid from tissues and returning it to the bloodstream. Because both systems work together to maintain fluid balance, understanding how dengue affects lymphatic vessels is important for explaining why fluid accumulation becomes so severe in critical disease.

In our study, we examined whether the viral protein NS1, which circulates during infection, directly affects the cells that line lymphatic vessels. We found that NS1 increases the permeability of these cells and reduces their ability to form new vessel structures. These effects were not caused by cell death but by disruptions in how the cells organise their junctions, internal scaffolding, and interactions with neighbouring cells.

By showing that NS1 can directly impair lymphatic vessel function, our work identifies a previously overlooked mechanism that may contribute to fluid build-up in severe dengue and suggests new avenues for future therapeutic research.

## Introduction

Dengue virus (DENV, *Orthoflavivirus denguei*) infection is responsible for ∼390 million infections per year worldwide (1), and has been included among the ‘Ten threats to global health’ in 2019 by the World Health Organisation (2). Each year, up to 96 million individuals manifest the clinical symptoms of the disease, with the majority presenting with uncomplicated dengue fever (DF) and a smaller subset developing severe dengue disease, which encompasses dengue haemorrhagic fever (DHF) and dengue shock syndrome (DSS) (2).

Severe dengue is characterised by deep vascular dysfunction, leading to vascular leakage, interstitial fluid accumulation and subsequent hypovolaemic shock and potentially death. Examples of common signs of clinically relevant fluid accumulation are the presence of pleural effusion and ascites resulting in respiratory distress and abdominal pain (3).

The role of vascular endothelial dysfunction in severe dengue vascular hyperpermeability and leakage has been linked to endothelial apoptosis, inflammatory cytokine storm, and the effect of secreted DENV non-structural protein 1 (NS1) on endothelial barrier function (4). NS1 is a viral protein secreted by cells infected with DENV and is found in abundance in the blood of patients affected by the disease. NS1 is involved in viral replication, pathogenesis, and immune evasion, and is also considered to play a key role in vascular leakage by promoting the disruption of the endothelial glycocalyx, therefore reducing endothelial barrier capacity (5). Our own work also demonstrated that NS1 has a strong effect on pericytes, a mural cell population ensheathing the capillary blood vessels and responsible for the regulation of the vascular barrier (6). Indeed, NS1 treatment of the co-culture of endothelial cells and pericytes, which more closely represent the *in vivo* vascular structure, resulted in a magnification of NS1-induced hyperpermeability (6).

Importantly, in physiological conditions, vascular extravasate is drained through the lymphatic vascular system to re-establish homeostasis and prevent an increase in interstitial pressure and fluid accumulation.

In the tissues, the lymphatic vasculature is composed of a blind-ended network of capillary vessels intermingled with blood vessels which drains the excess interstitial fluid, macromolecules and immune cells through large collecting lymphatic ducts into the lymph nodes before ultimately returning them to the blood circulation (7). Lymphatic vessels are formed of a monolayer of lymphatic endothelial cells displaying discreet permeability properties along the vascular tree; lymphatic capillary endothelial cells form ‘button’ junctions that act as one-way valves that are designed to favour the entry of the interstitial fluid into the lymphatic system when the pressure increases. Further down, lymphatic endothelial cells of the collecting vessels form more impermeable ‘zipper’ junctions, reminiscent of the blood vessel endothelium (7).

Lymphatic vessels’ role in maintaining interstitial fluid balance and returning extravasated liquid into the blood circulation is crucial for tissue homeostasis. The lymphatic system also participates in the immune response by recirculating immune cells, a system that is leveraged by dengue virus both to enter the blood circulation and to reach target tissues (8–10). Lymphatic hyperpermeability, leading to lymph extravasation is a reported issue in diseases affecting the vasculature, such as diabetes (11). Similarly to blood vessels, effective lymphatic function is dependent on tight control of the lymphatic endothelial layer permeability, and loss of lymphatic barrier function may represent an unexplored contributor to severe dengue pathology.

Severe dengue symptoms are compatible with a dysfunction of the lymphatic system, alongside the better characterised microvascular dysfunction, nevertheless studies on the effect of DENV and DENV NS1 on lymphatics are scarce.

In this study, we now show that DENV-2 NS1 also affects lymphatic endothelial cell permeability through the disruption of the junctions between lymphatic endothelial cells. Dengue-dependent lymphatic dysfunction and leakage, here described for the first time, is therefore a likely contributor to the haemorrhagic manifestations and fluid accumulation occurring in severe dengue disease.

## Materials and Methods

### Cell culture

Primary Human Dermal Lymphatic Endothelial Cells (HDLEC, C-12216) from juvenile foreskin and human umbilical vein endothelial cells (HUVECs, C-12203) from pooled donors were obtained from Promocell (Heidelberg, Germany). HDLECs were cultured Endothelial Cell Growth Medium MV 2 (EGM-MV2, #C-22022) and HUVECs in Endothelial Growth Medium 2 (EGM-2) (Promocell) in an incubator set at 37⁰C with 5% CO2.

Upon reaching 80% confluency, cells were subjected to trypsinization using DetachKit (PromoCell), according to the manufacturer’s instructions. For all experiments, HDLECs were used between passages 5 to 9. Recombinant DENV-2 NS1 PROTEIN (HEK293) from Native Antigen (DENV2-NS1-100 in DPBS) was used at a concentration of 1 µg/ml.

### Lymphangiogenesis assay

96-well plates were coated with 45 μl per well of cold Geltrex (Fisher Scientific cat# A4000046703, Growth Factor Reduced) and incubated for 30 min at 37°C to allow gelation. HDLECs were seeded at 1.2 x 10^4^ cells per well on the solidified Matrigel in 200 µl of EGM-MV2 in the presence or absence of 1 µg/ml DENV-2 NS1. The cells were cultured for 6 h in a 5% CO_2_ incubator at 37°C. Four or five images per well were acquired at 4X magnification using a Nikon Eclipse TS2 microscope (Nikon, The Netherlands). Images were analysed using ImageJ 1.53c (https://imagej.net/ij/). Six independent experiments were conducted, each with four technical replicates. Results were normalised to the mean of the untreated controls.

### Immunofluorescence staining

HDLECs were seeded on glass coverslips previously coated with 15 μg/ml fibronectin solution (P8248, Innoport) and cultured to confluency. Cells were washed with PBS and fixed with 4% paraformaldehyde (PFA) for 10 min at room temperature and then permeabilized with 0.02% Triton-X100 for 10 minutes and blocked with 10% goat serum (G9023- Merck) in PBS for 30 min. After blocking, the cells were incubated at 4°C overnight with the primary antibodies: anti-VE-cadherin (1:100, cat# sc-9989, Santa Cruz Biotechnology, Dallas, TX, USA), anti-ZO-1 (1:150, ZO-1 Polyclonal antibody 21773-1-AP, Proteintech: Manchester, UK), or anti-claudin 5 (1:50, rabbit anti-claudin-5, Fisher Scientific, cat# 341600). After overnight incubation, primary antibodies were washed with PBS and the appropriate secondary antibodies were added for 1 h at room temperature in the dark: anti-mouse and anti-rat Alexa Fluor-488 secondary antibody (1:500, ThermoFisher Scientific, Carlsbad, USA). Directly conjugated F-actin (Phalloidin-iFluor 555, 1:2000, cat# ab176756; Abcam, Cambridge, UK) was also incubated for 1 h at room temperature in the dark. The slides were then incubated with the nuclear stain DAPI (cat# D9542, Merck) at 1:1000 dilution for 10 min and mounted onto glass slides using mounting medium (Invitrogen™ Fluoromount-G™ Mounting Medium-00495802). The samples were imaged using a laser confocal microscope (ZEISS LSM 980 with Airyscan 2; Carl Zeiss, Jena, Germany).

### Image analysis

Image analysis and quantification was carried out using ImageJ-Fiji software (ImageJ 1.53c (https://imagej.net/ij/). Where brightness and contrast adjustments were necessary, they were applied to all images equally. Biological and technical replicates are indicated in figure captions.

Relative protein fluorescence intensity was quantified by measuring the integrated density of each image and subtracting background signal to obtain the corrected total cell fluorescence (CTCF). CTCF values were normalized to the number of nuclei per field and subsequently normalized to the untreated condition.

VE-cadherin junction width was quantified by manual measurement of cell–cell junctions at multiple regions within each field of view across independent biological replicates.

### Permeability assay

Transendothelial electrical resistance (TEER) assay was performed to determine HDLECs permeability. Firstly, inserts were coated with 15 μg/ml fibronectin solution and transferred to a 24 multi-well plate. Each insert was seeded with 5 x 10^4^ HDLEC in 200µL of fresh medium. The day after, medium was changed, and TEER was measured by an epithelial volt/ohm manual meter (EVOM™) combined with STX4 electrode. Three inserts with just EGM-2 medium but no cells were used as a control. After an additional 24 h; 10% of the media was replaced with media-containing DENV-2 NS1 or fresh media only. Permeability measurements were made at 0 h, 14 h, 16 h, 18 h, 20 h, 22 h and 24 h. The experiment was repeated in three independent biological replicates with three wells per condition and two measurements per well. Three inserts with just EGM-MV2 medium but no cells were used as a control. Raw resistance values (Ω) were obtained from cell-containing inserts and corresponding blank inserts (cell-free). To account for background resistance, blank values were subtracted from each measurement. The resulting values were multiplied by the membrane surface area (0.3 cm²) to obtain TEER values expressed in Ω·cm². For time-course analyses, TEER values were normalized to untreated controls at each time point. For percentage change analyses, TEER values were normalized to baseline (0 h) within each replicate using the following formula: ΔTEER (%) = (TEER₍24h₎ − TEER₍0h₎) / TEER₍0h₎ × 100. Data are presented as mean ± SD of independent replicates.

### Live/dead assay

HDLECs were seeded in 96-well plates at a density of 5 x 10^3^ cells/well and incubated overnight at 37°C. Cells were stained with calcein AM (10 mM) and EtBr (2 mM) for 30 min at 37°C. The positive control for the EtBr staining was fixed with 4% paraformaldehyde for 15 min and stained as described above. Cells were imaged using a Nikon Eclipse TS2 microscope. Two images per well were acquired at 4X and 10X magnification. ImageJ software was used to quantify calcein fluorescence and results were normalised to untreated controls. The results were obtained from three independent experiments with 3–4 technical replicates.

### MTT assay

An MTT assay was performed to further evaluate the viability and proliferation of HDLECs. The cells were seeded at a density of 1.2 x 10^4^ cells per well, in quintuplicate. Following a 24 h incubation at 37°C and 5% CO_2_, half of the wells were treated with DENV-2 NS1. After another 24 h incubation, MTT reagent 0.5 µg/mL was added and incubated for 4 hours, forming formazan crystals, which were then dissolved in sodium dodecyl sulfate and hydrochloric acid. After 15 minutes agitation and overnight incubation at 37°C, absorbance at 570 nm was measured using microplate reader (SpectraMax® iD3-5621, Molecular Devices, San Jose, CA, USA). to determine cell viability, with four independent experiments conducted.

### Wound healing assay

HDLECs were seeded 2 × 10^5^ cells per well in a 48-well plate and cultured to achieved complete confluency. A 200 ml pipette tip was used to create a scratch-wound in the cell monolayer. The wells were then washed twice with medium, followed by the addition of DENV-2 NS1 or control medium. Images of the scratch area were captured every 2 h for 48 h using a Cytation 5 WFOV (Agilent BioTek) Cell Imaging Multimode Reader. Cells were maintained between imaging intervals in a 5% CO_2_ atmosphere at 37°C. Three independent experiments were conducted, with three technical replicates. Images were analysed to calculate the wound size over time using ImageJ ‘Wound healing size’ plugin tool. Results are presented as the area percentage correlated to the initial wound size of 100%.

### RNA-Sequencing and analysis

HDLECs were seeded at a density of 1-3 × 10^5^ cells per well in 6 well plates, cultured to confluency and then treated with NS1 for 24 hours. Following treatment, total RNA was extracted using the Direct-zol RNA Microprep Kits (Zymo Research, cat# R2060) following the manufacturer’s instructions.

Bulk mRNA sequencing was performed by Novogene (UK) Company Limited. The experimental procedure for transcriptome sequencing includes sample quality control using Agilent 5400, library preparation using a kit with poly-T oligo-attached magnetic beads, cDNA synthesis, and quality control using Qubit and real-time PCR. Sequencing using Illumina technology with a “Sequencing by Synthesis” method.

The bioinformatics analysis pipeline involves data quality control with fastp software, reads mapping with HISAT2 (v 2.2.1), and gene expression quantification using featureCounts (v 2.0.6). Differential expression analysis is conducted with DESeq2 (v 1.42.0) or edgeR (version 4.0.16), and enrichment analyses are performed using clusterProfiler (v 4.8.1). Additional analyses include GSEA, PPI analysis via STRING, alternative splicing with rMATS (v 4.3.0), SNP analysis using GATK (v 4.1.1.0), and fusion gene detection with STAR. Raw RNA-Seq gene counts were batch-corrected using ComBat-seq (12), and differential gene expression analysis comparing NS1 treatment against vehicle control was performed using DESeq2 (N = 4) (13). Principal component analysis (PCA) was performed in R on batch-corrected data and plotted in GraphPad Prism version 10.4.1. Differentially expressed genes (DEGs) were submitted to DAVID (14) for functional enrichment analysis to identify significantly enriched KEGG pathway terms following NS1 treatment. Significant terms were plotted using ggplot in R. Significantly DEGs and enriched KEGG pathways were adjusted for multiple hypothesis testing using the Benjamini and Hochberg method (padj < 0.05).

### Microfluidic experiment

Blood vessel and lymphatic endothelial cells were seeded on an OrganoPlate® 3-lane (MIMETAS, 4004-400-B) chip separated by a central channel filled with a gel consisting of 2 µL of 5 mg/ml collagen-1 (Corning® Collagen I, cat# CLS354236) neutralised with 10% Na_2_CO_3_ (37 g/l, pH 9.5) and 10% HEPES buffer (1 M). The plate was incubated at 37°C for 15 minutes to polymerise the collagen gel. HDLEC and HUVEC cells were suspended in their respective media at 10,000 cells/µL, and 2 µl of cell suspension dispensed into the inlet of one of the perfusion channels.

Media (50 μl) was loaded into the perfusion outlet. The plate was placed on its side at an angle of 75° for 3-4 h in a 37°C incubator to allow the cells in the channels to settle and attach. 50 μl of media was added to the perfusion inlets. The plate was incubated at 37°C on a rocker platform exposed to a bi-directional flow generated by passive levelling of reservoirs through gentle, gravity-driven rocking. The cells formed a monolayer within 24-48 hours, and the media was changed every other day for 14 days.

### Statistical analysis of in vitro assays

Statistical analyses were performed using Prism software version 10.0 (GraphPad Software, San Diego, CA, USA). The data passed the D’Agostino-Pearson omnibus normality test. Data are presented as mean ± standard deviation (SD), and statistical significance was set at p < 0.05. Comparisons between two groups were assessed using two-way unpaired Student’s t-test. Comparisons involving more than two groups or multiple factors were assessed using one or two-way ANOVA followed by Bonferroni post-test for individual comparisons.

### Data Availability Statement

The RNA-Seq data generated in the present study have been deposited to the NCBI Gene Expression Omnibus (GEO) with the GEO accession [GSE326043].

## Results

### Lymphatic endothelial cell permeability is compromised by DENV-2 NS1 treatment

DENV circulating protein NS1 is known to cause endothelial dysfunction in blood vessels (5) but despite the importance of the lymphatic system in clearing accumulated excess interstitial fluid and cells, its effect on lymphatic endothelial cells has not yet been reported.

To test the effect of DENV-2 NS1 on the lymphatic barrier function, we seeded primary human dermal lymphatic endothelial cells (HDLEC) on transwells and cultured them to confluency. Cells were then treated for 24 h with 1μg/ml DENV-2 NS1 or control medium, and transendothelial electrical resistance (TEER) was measured at regular intervals to establish changes in the permeability of the lymphatic endothelial cell layer (Figure 1A).

**Figure 1.**
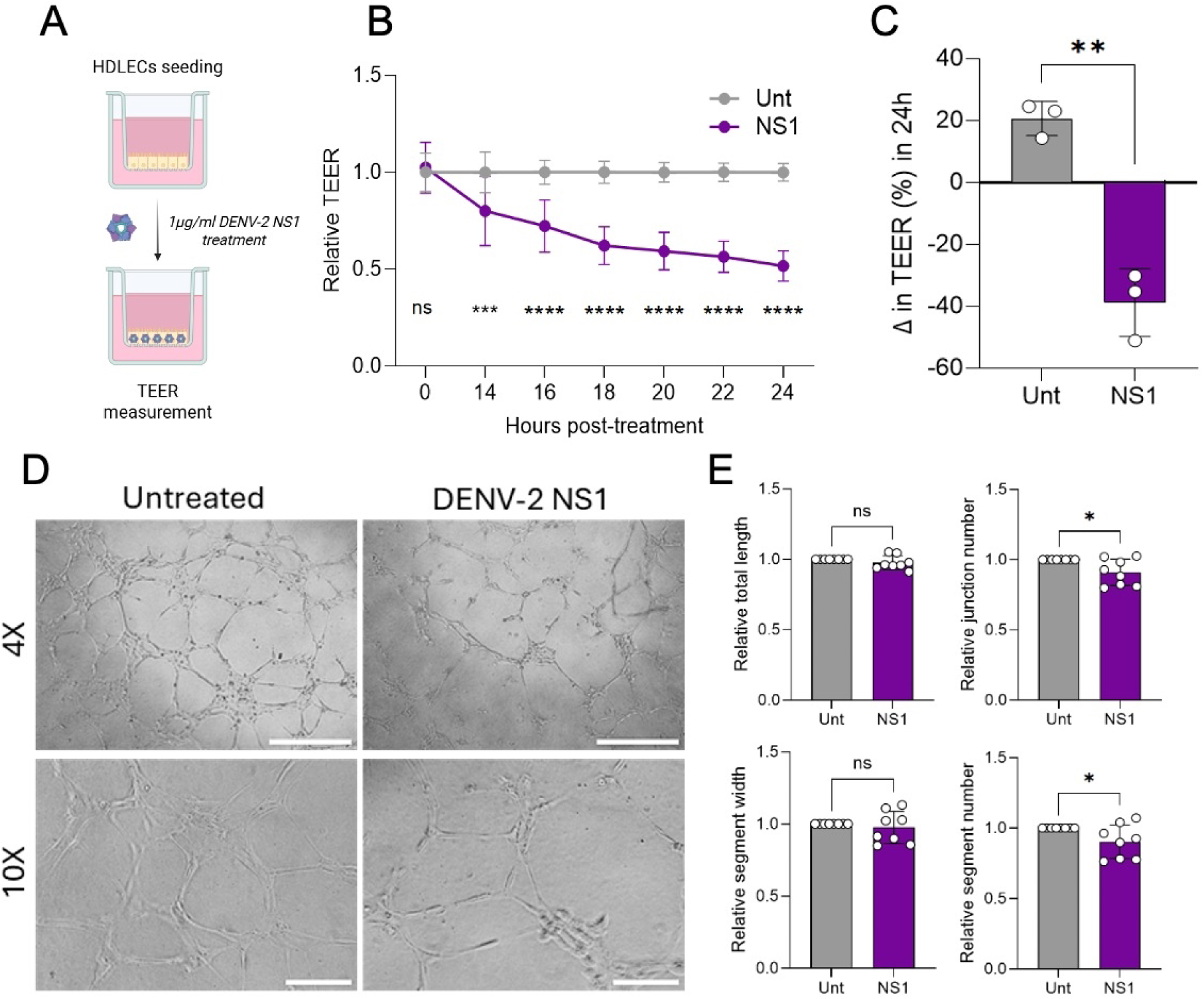
DENV-2 NS1 decreases HDLECs barrier function and angiogenic capacity. (A) Schematic for transendothelial electrical resistance assay set up (TEER). (B) Change in TEER values over 24 h in untreated and NS1-treated HDLECs. N = 3, n = 3. Statistical significance was determined by two-way ANOVA with Bonferroni’s multiple comparisons test. (C) Overall change in TEER over the 24 h period as a percentage. N = 3, n = 3. Statistical significance determined by unpaired two-tailed t-test). HDLECs grown on angiogenic matrix in the presence or absence of 1 μg/mL DENV-2 NS1. (D) Representative images of untreated and NS1-treated HDLECs tube-like structures. Scale bar: 250 μm (4x); 50μm (10x). (E) Quantitative analysis of tube-forming assay using ImageJ Vascular Analyzer; “total network length” calculated as total length of structures within field of view, “number of junctions” defined as connecting vessel-like tubular structures, “number of segments” and “segment width” calculated as the total number and the mean width of all structures connected to junctions at both ends. N = 6, n = 4. Statistical significance was established by unpaired two-tailed t-test. Results are shown as mean ± SD, ns= non-significant, *p ≤ 0.05, ***p ≤ 0.001 ****p ≤ 0.0001. Unt, untreated. NS1: DENV-2 NS1.

The results showed that HDLEC treated with DENV-2 NS1 progressively lose their barrier function over the 24h period, leading to almost 50% increase in permeability, with no recovery measured over this period (Figure 1B). The maximum change in TEER was overall strongly negative exclusively in cells treated with DENV-2 NS1 (Figure 1C).

These results show for the first time that DENV NS1 not only affects the permeability of capillary-derived endothelial cells, but also affects lymphatic permeability.

### DENV-2 NS1 treatment reduces lymphangiogenesis

Maintaining lymphatic vessel density throughout the tissue is essential for effective fluid absorption. Lymphangiogenesis (formation and maturation of new vessels from existing ones) can occur at sites of inflammation, driven by pro-inflammatory molecules released by macrophages and granulocytes, and facilitates the resolution of tissue oedema and enhances immune responses by promoting macrophage and dendritic cell mobilization (15).

To assess the effect of DENV-2 NS1 on lymphangiogenesis, HDLEC were cultured on the 3D angiogenic matrix Matrigel and the organisation of the cells into vessel-like structures was quantified at 6 h post-seeding in the presence or absence of DENV-2 NS1. HDLEC efficiently organised into tube-like structures on the matrix when cultured in control medium; conversely, the structures formed in the presence of NS1 appeared defective and sparser (Figure 1D). Indeed, quantification of the vessel-like features showed that DENV-2 NS1 significantly reduced the number of junctions and number of segments, structures connecting two junctions (Figure 1E). No statistical difference was detected in the total length of the structures and in their thickness (Figure 1E). These data suggest that DENV-2 NS1 interferes with the HDLECs intrinsic lymphangiogenic capacity and in particular reduces their ability to form complex and interconnected networks.

### Lymphatic endothelial cell viability, size and migratory capacity are unaffected by DENV-2 NS1 treatment

The reduction in lymphatic endothelial cell barrier function and lymphangiogenic potential could derive frim reduced cell numbers due to lower proliferation or increased cell death. To rule out such nonspecific effects of NS1, cell viability was measured using the metabolic MTT assay. Our results indicated no difference in the quantitative fluorescence output of the assay in NS1-treated HDLECs compared to control treatment, suggesting that the overall viability and cell number was not affected by DENV-2 NS1 treatment (Figure 2A). To quantify cell death in our cultures, HDLEC were treated with NS1 for 24 h or left untreated, then stained with Calcein-AM, a cell-permeant dye that is processed into a fluorescent molecule inside live cells (live staining), and Ethidium homodimer-1, a fluorescent molecule with high affinity for nucleic acids which only permeates dead cells (dead staining). Our results showed that in both control and NS1-treated HDLEC, cells were stained uniformly positive for Calcein-AM with no presence of apoptotic Ethidium homodimer-1 positive nuclei, indicating that NS1 treatment does not affect lymphatic endothelial cell viability (Figure 2B and C).

**Figure 2.**
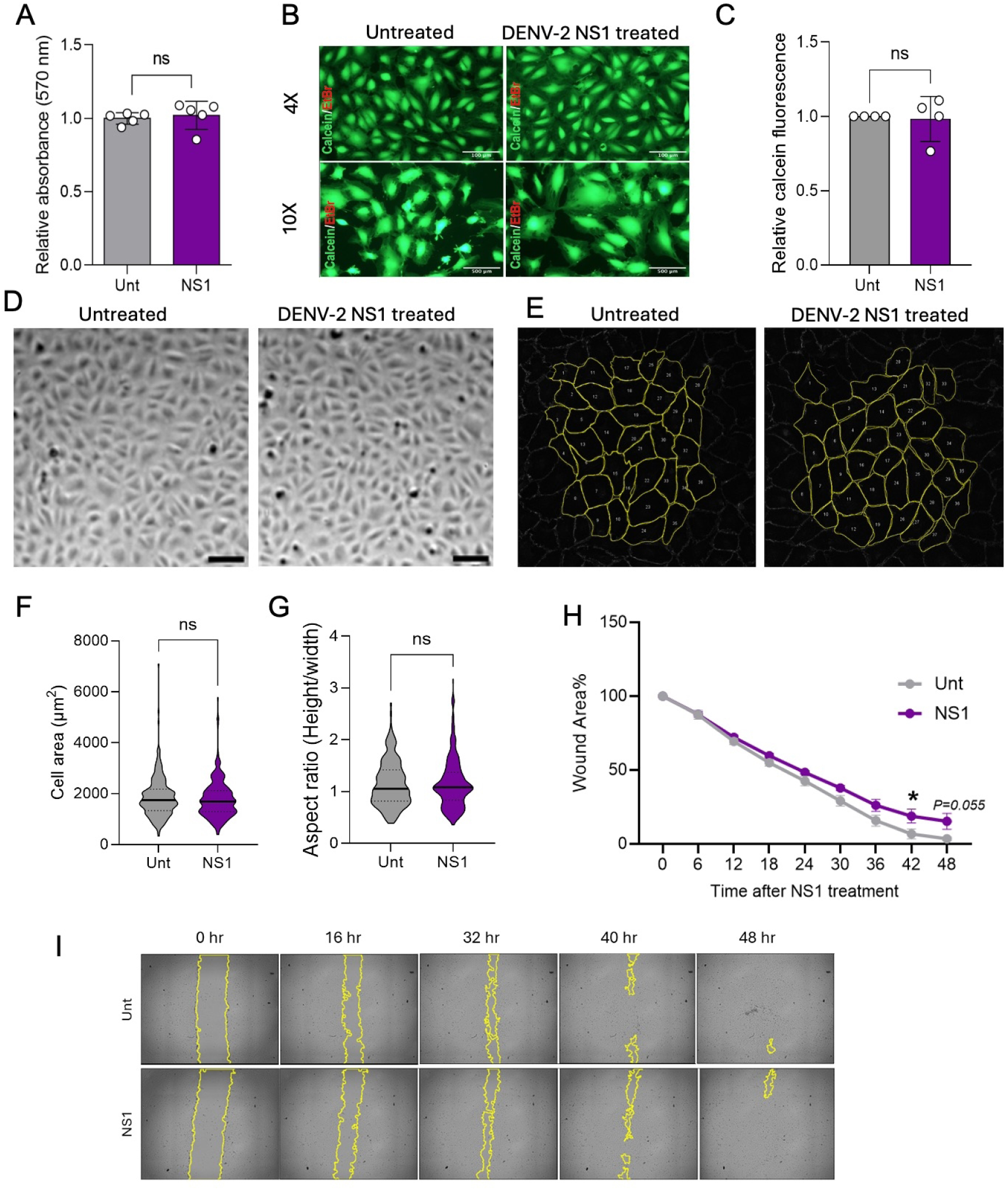
DENV-2 NS1 does not affect HDLECs viability, shape or size but reduces migration capacity. (A) HDLEC viability in response to 1 μg/mL DENV-2 NS1 treatment (NS1) assessed by MTT assay and normalised to untreated controls (Unt), and (B) by live/dead assay as shown in representative images of calcein (green, live) and ethidium bromide (red, dead) stained cells and (C) relative quantification of green fluorescence. N = 4-5, n = 5. Statistical significance was determined by unpaired two-tailed t-test. Scale bar: 150 μm (4x); 50μm (10x). (D) Representative bright field images of HDLEC untreated or treated with 1 μg/mL DENV-2 NS1 and (E) representative skeletonised images used to quantify the (F) cell area and (G) morphology using aspect ratio, aspect ratio = width/height. N=3, analysing 160-190 cells per experiment. Statistical analysis was performed using unpaired-sample two-tailed t-tests. Wound healing assay was performed on confluent HDLECs treated with1 μg/mL DENV-2 NS1 or control medium. (H) Analysis of the wound size reduction in percentage was performed to quantify migratory capacity. (I) Representative images show the progression of wound closure over time (I). N=3, n = 3. Results are shown as mean ± SD. Unt, untreated. NS1: DENV-2 NS1.

Cell shape and size changes can also be indicative of a process of differentiation or activation of endothelial cells, potentially contributing to loss of barrier function. However, brightfield images of untreated and NS1-treated cells did not display overt difference in cell morphology (Figure 2D), which was reflected in the quantification of average cell area and aspect ratio in skeletonised images (Figure 2E-G).

Endothelial cell migration is crucial for the development of new vessels and the maintenance of a coherent cell monolayer upon external challenge. To test the effect of NS1 on cell migration, HDLECs were seeded to confluency and the monolayer was wounded using a pipette tip to form a homogenous interruption in the cell monolayer. This process stimulates the cells to migrate and ‘repair’ the wound; the speed and efficiency of closure can be measured using longitudinal imaging. Both control and NS1-treated monolayers were capable of closing the wound almost completely over 48 h; however, NS1 treatment induced a delay in wound closure (Figure 2H-I), indicating a defect in cell migration.

Taken together, our data show that NS1 treatment is not affecting lymphatic endothelial cell proliferation, viability, or size. However, NS1 does reduce cell migration, which may contribute to the increased barrier permeability observed in Figure 1.

### Lymphatic endothelial cell-cell junctions and cell-matrix interactions are compromised by NS1 treatment

To explore potential molecular mechanisms underpinning the dysfunctional behaviour of lymphatic endothelial cells following exposure to NS1, we performed transcriptomic profiling analysis on HDLECs treated with DENV-2 NS1 for 24 h and compared gene expression with control medium-treated cells. The results showed a good separation of the two experimental groups as shown by the principle component analysis plot (Figure 3A). Bioinformatic analysis of the differentially expressed genes (DEGs) between NS1- and control-treated cells revealed a statistically significant (padj <0.05) enrichment of four pathways from the Kyoto Encyclopedia of Genes and Genomes (KEGG) database related to ‘focal adhesion’, ‘regulation of actin cytoskeleton’, ‘tight junction’ and ‘protein processing in endoplasmic reticulum’ (Figure 3B). While most of the DEGs were unique to one the of KEGG pathways, several contributed to more than one pathway (Figure 3C). Importantly, these pathways are central to both lymphangiogenesis and barrier maintenance in lymphatic vessels(7).

**Figure 3.**
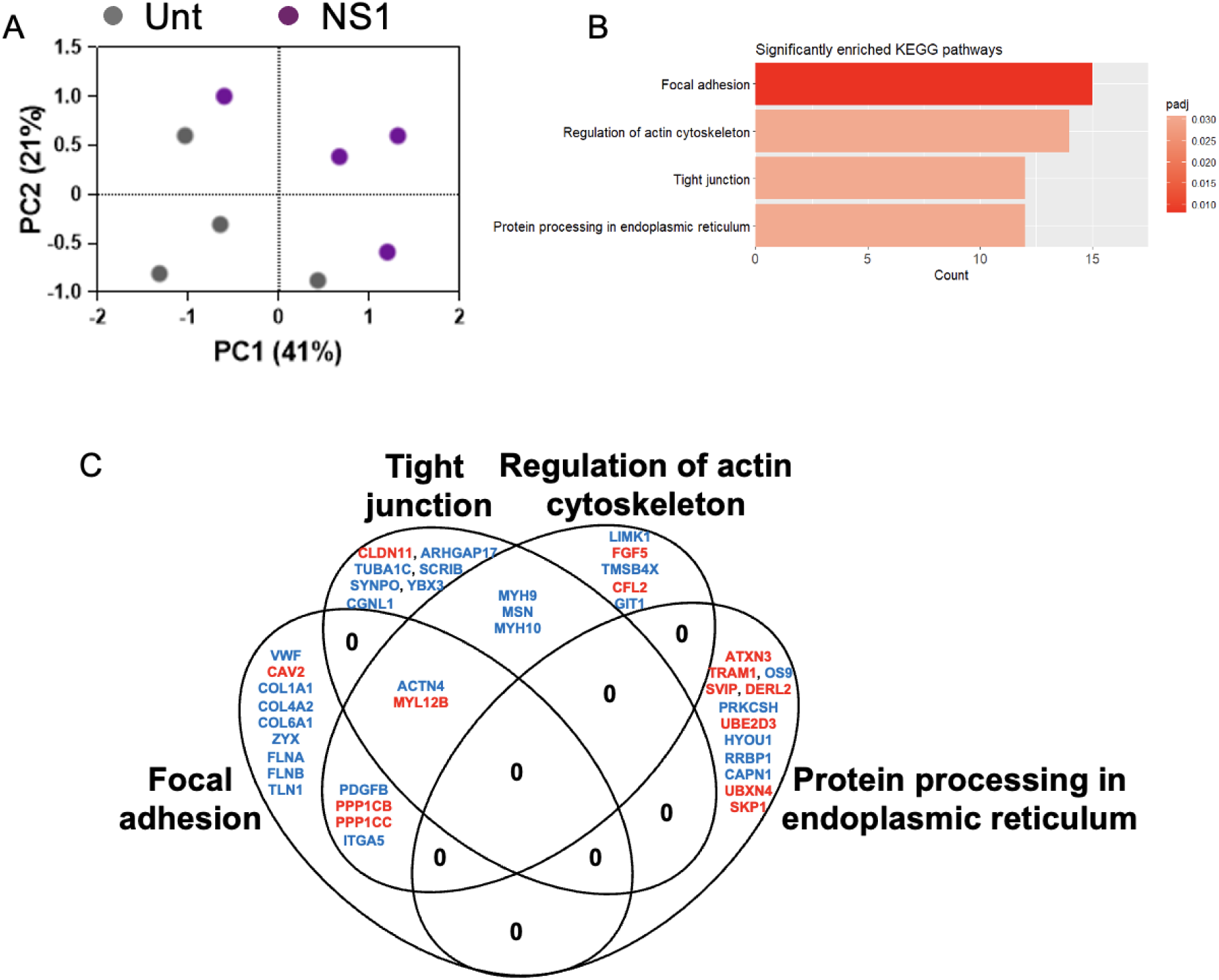
Transcriptomics analysis reveals NS1-dependent changes in cell-cell and cell-matrix adhesion pathways. Principal Component Analysis (PCA) plot of control (untreated=Unt) and DENV-2 NS1-treated (NS1) HDLECs (N = 4) (A). Bar plot of KEGG pathway terms significantly enriched in DENV-2 NS1-treated HDLECs, relative to control (B). For DEGs and enriched terms, significance values are corrected for multiple testing using the Benjamini and Hochberg method (padj < 0.05). Venn diagram illustrating the contribution of DEGs to the different KEGG pathways identified (C; blue=upregulated and red=downregulated).

Tight junctions mediate cell-to-cell adhesion, forming a sealing barrier between endothelial cells to prevent leakage. Several genes involved in tight junctions were differentially expressed upon NS1 treatment: claudin-11 (CLDN-11), Rho GTPase-activating protein 17 (ARHGAP17), tubulin alpha-1c chain (TUBA1C), Scribble (SCRIB), Synaptopodin (SYNPO), Y-box-binding protein 3 (YBX3), and cingulin-like 1 (Cgnl1). The tight junction proteins of the claudin and occludin families can also associate with the cytoskeleton through zona occludens (ZO) proteins, and in line with this, some DEGs in this KEGG are shared with ‘regulation of actin cytoskeleton’, including: myosin-9 and 10 (MYH9 and MYH10) and moesin. Others are shared additionally with ‘focal adhesion’: including alpha-actinin-4 (ACTN4) and Myosin Light Chain 12B (MYL12).

Focal adhesions mediate the anchoring of the endothelial cells to the extracellular matrix, NS1 induced DEGs uniquely associated with this KEGG pathway: including the cell adhesion protein zyxin (ZYX) and talin-1 (TLN1), cell filament components (FLNA and B), secreted protein von Willebrand factor (vWF), caveole-associated caveolin-2 (CAV2) and extracellular matrix components such as collagens (COL1A1, COL4A2 and COL6A1). Given the close association of focal adhesions with the cytoskeleton, several DEGs were shared between these two KEGGs: platelet-derived growth factor B (PDGFB), beta and gamma catalytic subunit of protein phosphatase 1 (PP1CB and PP1CB), and integrin alpha V (ITGA5).

Additionally, several DEGs uniquely clustered in the ‘protein processing in endoplasmic reticulum’ group, encoding for genes involved in protein degradation, autophagy and membrane trafficking: ataxin-3 (ATXN3), ubiquitin conjugating enzyme E2 D3 (UBE2D3), endoplasmic reticulum lectin 2 (OS9), small VCP interacting protein (SVIP), derlin-2 (DERL2), UBX domain-containing protein 4 (UBXN4), S-phase kinase-associated protein 1 (SKP1), autophagy-associated proteases calpain-1(CAPN1), protein transportation through the endoplasmic reticulum (ER), such as translocating chain-associated membrane protein 1 (TRAM1), protein folding and quality control beta-subunit of the enzyme glucosidase II (PRKCSH), hypoxia up-regulated 1 (HYOU1) and ribosome binding to the ER, ribosome-binding protein 1 (RRBP1).

The results from the RNAseq analysis therefore indicate that lymphatic endothelial cells exposed to DENV-2 NS1 undergo significant remodelling of cell-cell and cell-matrix junctions which may help explain the functional changes in lymphangiogenesis and permeability we observed.

### DENV-2 NS1 induces a profound remodelling of lymphatic endothelial cell junction proteins

The results obtained from transcriptomics analysis suggest that NS1 causes disregulation of lymphatic endothelial cell junctions. Endothelial cell barrier function and angiogenic potential are highly dependent on reciprocal cell adhesion and the anchoring of cells to the extracellular matrix, ensuring the maintenance of a selectively permeable barrier between the intravascular and the interstitial spaces and the correct organisation of vascular structures. Vascular endothelial (VE)-cadherin is an endothelial cell specific adhesion molecule that plays a crucial role in cell-cell junctions and regulation of vascular permeability and sprouting HDLEC were treated with NS1 for 24 h and then stained for VE-cadherin (Figure 4A). Quantification of VE-cadherin intensity showed an increase in VE-cadherin intensity in treated cells (Figure 4B). Correspondingly, VE-cadherin junctions were wider, deviating from the classical linear pattern of functional junctions found in controls (Figure 4C).

**Figure 4.**
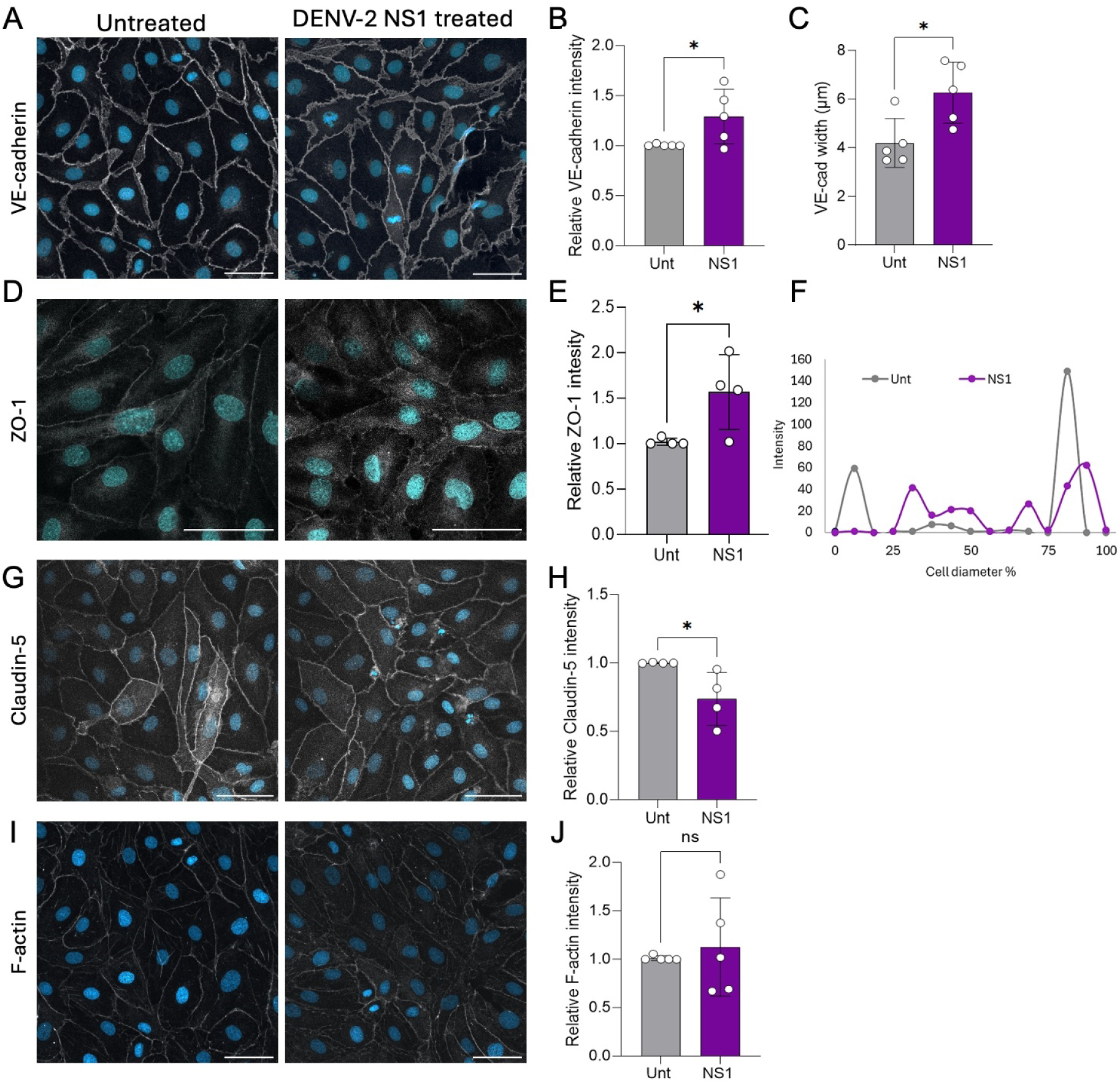
Junctional protein organisation is affected by DENV-2 NS1 treatment. HDLEC were treated with 1 μg/mL DENV-2 NS1 (NS1) or control medium (untreated=Unt) for 24 h and then stained by immunocytochemistry. (A) Representative images, (B) quantification and (C) junction width analysis of HLDEC monolayers stained for VE-cadherin (white) Scale bar: 50µm. N=5 (B and C), analysing 100–600 junctions from each replicate (C). (D) Representative images, (E) quantification of ZO-1-stained HLDECs (white) and (F) representative intensity profile across one individual cell. Scale bar: 50µm. N = 4. (G) Representative images and (H) quantification of Claudin-5 (white). Scale bar: 50µm. N = 4. (I) Representative images and (J) quantification of F-actin (white). N = 4-5. Nuclei are stained with DAPI, blue. Results are shown as mean ± SD. All samples analysed for statistical significance using unpaired-sample two-sided t-tests. ns=not significant *p ≤ 0.05, **p ≤ 0.01.

HDLEC were also stained for claudin-5 and ZO-1, two prominent components of the tight junction machinery in lymphatic cells(16). Similarly to VE-cadherin, ZO-1 staining showed an overall increase in fluorescence (Figure 4D and E). Further analysis revealed a redistribution of the protein from the cell membrane to the cytoplasm of cells treated with NS1, resulting in an overall apparent increase in intensity as confirmed by single cell fluorescence profiling (Figure 4F). Claudin-5 staining showed disrupted areas of discontinued junctions, reflected in reduced overall fluorescence (Figure 4G and H).F-actin staining was used to identify actin filaments and demonstrated that NS1 elicited changes in the organisation of the cytoskeleton albeit not reflected in a change of the total amount of fluorescence (Figure 4I and J).

Overall, these data confirm changes in the cytoskeleton and cell junctions identified by RNAseq analysis following DENV-2 NS1 treatment.

### NS1 disrupts lymphatic-like vessels 3D culture

Lymphatic vessels are embedded within organ tissues, often in very close contact with blood vessels. Luminal flow is a force that notably shapes the physiology of all endothelial cells, with blood vessel endothelial cells exposed to higher levels of shear and lymphatic endothelium experiencing a more sluggish flow. Recent work has highlighted the importance of flow on lymphatic endothelium physiology, showing an increase of cell alignment and junctional organisation under physiological conditions (17,18). We integrated low-shear stress and proximity culture with blood vessels in a single model using Mimetas 3-lane organoplates by seeding human umbilical vein endothelial cells (HUVECs) on the top channel and HDLECs in the bottom channel, separated by a collagen gel in the middle (Figure 5A). The plate was placed on its side for 3-4 h to allow cells to settle and then flow was applied by culturing the plate on a rocker shaker for 14 days. During this time period, 3D vessel wall-like structures formed in both blood vessel and lymphatic endothelial channels (Figure 5B and C). Staining and confocal imaging revealed that the co-culture conditions in the organoplate established strong cell-cell junctions in HDLEC, as shown by the robust expression of VE-cadherin and organised distribution of the cytoskeleton (Figure 5D). Interestingly, treatment of HDLECs cultured in this system with DENV-2 NS1 for 24 h caused dysregulation of VE-cadherin junctions with loss of localisation at the junctions, and a disruption of the cytoskeleton organisation (Figure 5D).

**Figure 5.**
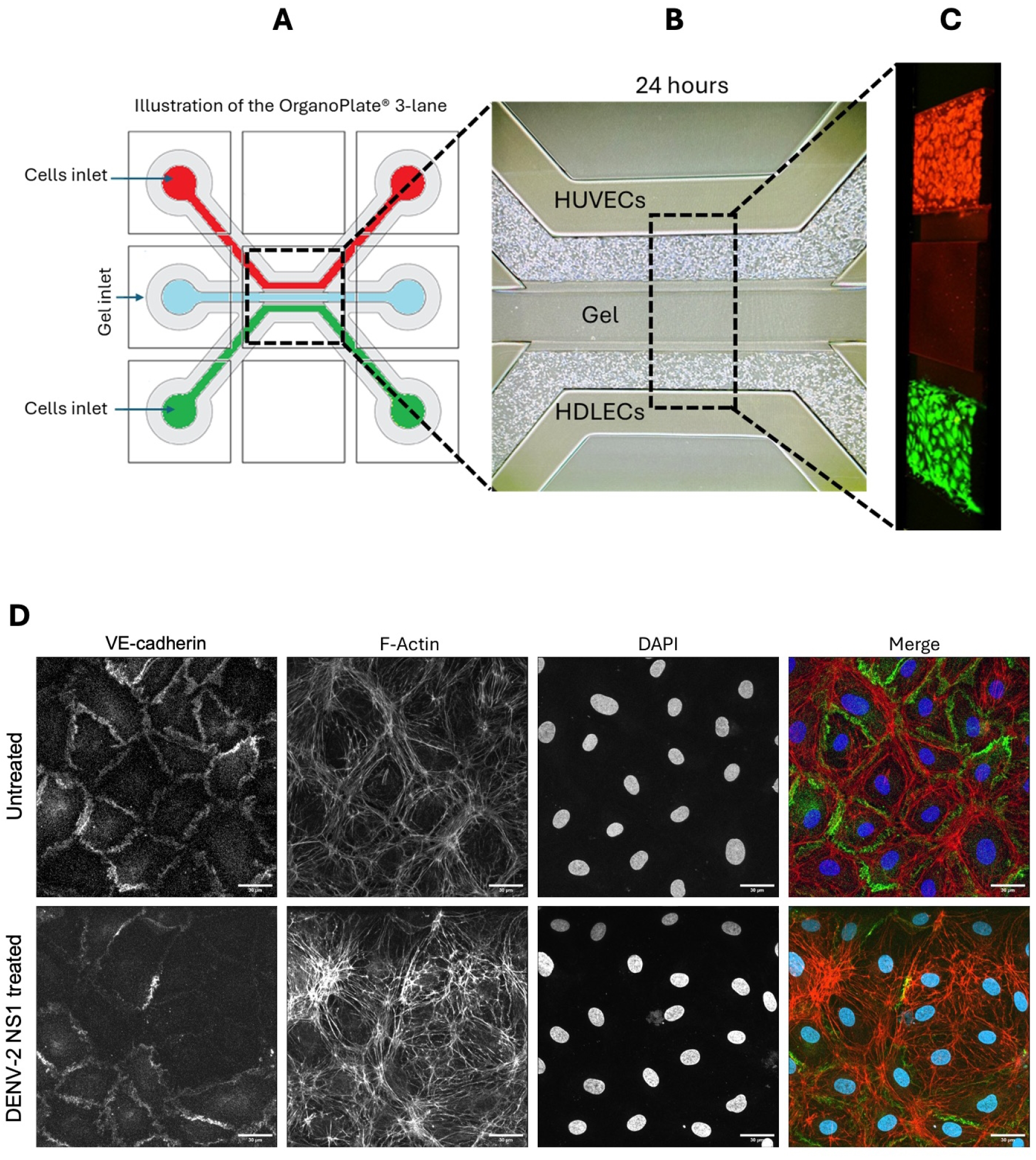
HDLECs cultured in flow conditions display profound changes upon DENV-2 NS1 treatment. (A) Schematic representation of an OrganoPlate® 3-lane tissue chip formed by two channels for cell seeding and a middle channel for gel loading. (B) Brightfield image and (C) confocal projection of HUVECs (red) seeded on top channel and HDLECs (green) seeded in bottom channel. Images taken after 24 h of culture under flow conditions. (D) Representative confocal max projection of HDLECs cultured in flow for 14 days in OrganoPlate® and treated for 24 h with DENV-2 NS1 (NS1) or control medium (untreated=Unt). Nuclei stained with DAPI. Merged images shows VE-cadherin (green) and F-actin (red) and DAPI (blue). Scale bar = 30 µm.

These data indicate that the culture of HDLECs in vessel-like structures under physiological conditions in the proximity of blood vessel endothelial cells confirms the disruption of lymphatic endothelium integrity following DENV-2 NS1 treatment, with a stronger emphasis on cytoskeleton remodelling.

## Discussion

The lymphatic system plays a crucial role in viral infections by coordinating the immune response and clearing accumulated interstitial fluid and cell debris. The lymphatic system can also be highjacked by viruses for dissemination throughout the body and to attract immune cells that are amenable to infection (19). We here show that DENV NS1 disrupts the lymphatic endothelial barrier in 2D and 3D cultures, including in vessels cultured under physiological conditions. Furthermore, these effects of NS1 were not linked to alterations in cell viability or proliferation, but were associated with an altered migration capacity of lymphatic endothelial cells and transcriptomic and cellular ultrastructure changes indicative of alterations in cell-cell junctions and the cytoskeleton.

### Lymphatic disruption during viral infection

The role of lymphatic vessels in dengue disease has so far only been reported in the context of a skin explant model, in which DENV infection of VEGFR3+ lymphatic endothelial cells was reported, albeit less inconsistently across specimens than for other cell types (20).

Since the skin mounts an antiviral immune response, dermal Langerhans cells (20) as well as circulating and tissue resident mononuclear phagocytes (MNP) and dendritic cells (DC) (9), which are key DENV target cells, become exposed to the virus and exports it to other susceptible tissues. Cell-free viral particles can also enter the lymphatic system and then infect lymph noderesident macrophages and T cells. Both cell-free virus and infected cell-mediated lymphatic dissemination routes have previously been described for closely related viruses such as West Nile virus and Zika virus (21).Additionally, given the relatively higher concentration of leukocytes and proteins in lymph compared to blood (22), it is also reasonable to expect a similar or higher concentration of NS1 in the lymph compared to blood, potentially leading to more pronounced impacts on lymphatic vessels.

Interestingly, work by us and others has previously reported different magnitudes of NS1 effects on endothelial permeability, largely dependent on the specific flavivirus and vascular cell origin (5,6,23–25). For DENV-2 NS1, a ∼20% decrease in barrier function in human blood vessel endothelial cells treated with 0.5-5 µg/ml DENV-2 NS1 from all tested organs as measured by TEER in accordance with the systemic nature of this disease (5,6,23,24). Resealts here presented showed that lymphatic endothelial cells displayed a more pronounced ∼50% reduction in barrier function upon stimulation with 1 μg/ml of NS1, suggesting a greater sensitivity of lymphatic endothelial cells to NS1.

### Dysregulation of lymphatic tight junctions by DENV NS1

Our results demonstrate that lymphatic endothelial cells respond to NS1 challenge by displaying global changes in genes involved in tight junctions, focal adhesions, and protein processing, results that are in line with previous work indicating deep remodelling of the cell-cell junction machinery in blood vessel endothelial cells that correlate with functional changes in permeability measured by TEER (5,26). Our immunocytochemical analyses confirmed that NS1-induced hyperpermeability was associated with a dysregulation of cell-cell junctions, as measured by VE-cadherin, claudin-5 and ZO-1 staining. VE-cadherin is a cadherin family protein that is primarily expressed in vascular endothelial cells and forms the core of adherens junctions (5). Intracellularly, VE-cadherin binds to actin filaments, through regulatory molecules such as catenins.

Previously, NS1 was shown to induce VE-cadherin internalisation via clathrin-mediated endocytosis in blood vessel endothelial cells, leading to reduction of expression at the membrane level (5). While we did not observe a decrease in VE-cadherin localisation on the cell membrane in lymphatic endothelial cells, we report a shift of the junction shape from ordered and linear to thick, heterogeneous and disorganised, compatible with a marked loss of barrier integrity (27). The central role of junction disruption in the NS1-dependent lymphatic barrier dysfunction is also confirmed by the RNA-seq data, which identified changes in tight junction and focal adhesion pathways and specifically pointed at the differential expression of several molecules that have been shown by proximity proteomics to interact with VE-cadherin on the membrane of lymphatic endothelial cells (28). This change in junction conformation is often associated with actin cytoskeleton remodelling, which pulls on the junctions radially, forming gaps (27). Indeed, our RNA-seq data highlighted changes in actin cytoskeleton regulation, also confirmed by immunostaining of cells, especially under physiological flow conditions.

Of note, the VE-cadherin junctions observed in our LDEC cultures exhibited ‘zipper’ junctions, replicating the impermeable endothelial layer of the collecing lymphaticscells reflect, larger vessels transporting the lymph to the lymph nodes, rather than the ‘button-like’ junctions of lymphatic capillaries. *In vivo*, such an increase in permeability in the collecting lymphatics would be expected to lead to a defect in the transport of interstitial fluid and inflammatory cells away from the affected tissues, exacerbating the effect of blood vessel haemorrhage and fluid extravasation (11). Furthermore, loss of integrity in collecting lymphatics may impair the fluid dynamics within the lymphatic system, reducing the suction created by lymphatic pumping (29). Importantly, diabetes, which is typically associated with blood vessel dysfunction, has also been shown to increase the permeability of ‘zipper’ junctions in collecting lymphatic vessels, potentially contributing to fluid accumulation in the tissues and oedema (11). Our results are the first to report a similar pattern of hyperpermeability in lymphatic cells in response to DENV NS1, associated with cell-cell junction remodelling. The synergistic failure of blood vessel and lymphatic barrier function may therefore contribute to the extravasate accumulation and hypovolaemic shock detected in severe dengue patients.

### Potential impacts of DENV NS1 on lymphangiogenesis

Lymphangiogenesis is a dynamic mechanism that regulates the number and density of lymphatic vessels in the tissue and is stimulated by increased demand after injury or by inflammatory mediators upon establishment of infection. Establishment of inflammation-induced lymphangiogenesis can hasten the resolution of diseases, providing increased capacity for liquid drainage (19). Our results showed a decrease in lymphangiogenesis in response to NS1, suggesting that NS1 is capable of inhibiting the formation of new lymphatic vessels, effectively blunting the host response to increased interstitial volume. Interestingly, the effect of lymphangiogenesis on viral resolution varies greatly depending on the virus and also the organ. For example, VEGF-A-stimulated lymphangiogenesis promotes herpes simplex virus type 1 (HSV-1) keratitis in the cornea (30), whereas influenza A virus (IAV) induces lung lymphangiogenesis that helps resolve lung oedema by enhancing drainage (31).

The formation of new blood vessels is intrinsically linked to endothelial cell viability, proliferation and migratory capacity (32). Our results indicated that NS1 treatment did not affect lymphatic endothelial cell viability or proliferation, which is in line with our previous findings in vascular cells (6). On the other hand, we identified a decrease in lymphatic endothelial cell migratory capacity, resulting in a delay in the wound closure which may explain the observed reduction in angiogenic capacity (33).

## Acknowledgements

We wish to thank Yan Zhang and Taija Mäkinen (Uppsala University, Sweden) for sharing their knowledge and expertise and providing insightful feedback.

## Data availability

The raw RNA-seq data generated during the current study have been deposited in the NCBI Gene Expression Omnibus (GEO) repository and are available under accession number GSE326043.

The datasets generated and analysed during the current study will be available upon publication in the Zenodo repository, DOI: 10.5281/zenodo.19105946.

